# CAR T cell infiltration and cytotoxic killing within the core of 3D breast cancer spheroids under control of antigen sensing in microwell arrays

**DOI:** 10.1101/2024.03.14.585033

**Authors:** Youngbin Cho, Matthew Laird, Teddi Bishop, Ruxuan Li, Elisa Ruffo, Jason Lohmueller, Ioannis K. Zervantonakis

## Abstract

The success of chimeric antigen receptor (CAR) T cells in blood cancers has intensified efforts to develop CAR T therapies for solid cancers. In the solid tumor microenvironment, CAR T cell trafficking and suppression of cytotoxic killing represent limiting factors for therapeutic efficacy. Here, we present a microwell platform to study CAR T cell interactions with 3D tumor spheroids and determine predictors of anti-tumor CAR T cell function. To precisely control antigen sensing by CAR T cells, we utilized a switchable adaptor CAR system, that instead of directly binding to an antigen of interest, covalently attaches to co-administered antibody adaptors that mediate tumor antigen recognition. Following addition of an anti-HER2 adaptor antibody, primary human CAR T cells exhibited higher infiltration and clustering compared to the no adaptor control. By tracking CAR T cell killing at the individual spheroid level, we showed the suppressive effects of spheroid size and identified the initial CAR T cell : spheroid area ratio as a predictor of cytotoxicity. Spatiotemporal analysis revealed lower CAR T cell numbers and cytotoxicity in the spheroid core compared to the periphery. Finally, increasing CAR T cell seeding density, resulted in higher CAR T cell infiltration and cancer cell elimination in the spheroid core. Our findings provide new quantitative insights into CAR T cell-mediated killing of HER2+ breast tumor cells. Given the miniaturized nature and live imaging capabilities, our microfabricated system holds promise for discovering cell-cell interaction mechanisms that orchestrate antitumor CAR T cell functions and screening cellular immunotherapies in 3D tumor models.

## Introduction

Chimeric antigen receptors (CARs) have emerged as a promising approach to develop cell-based immunotherapies^1-3^. CARs, composed of an antigen-specific antibody single chain variable fragment fused to T cell signaling domains, have been used to engineer T cells that are activated upon binding to a target antigen. CAR T cell-based immunotherapies have been FDA-approved in acute lymphoblastic leukemia and multiple myeloma^4-7^. Developing CAR T cells that recognize targets enriched in solid tumors, such as human epidermal growth factor receptor 2 (HER2) has been a major research focus, yet the clinical trials have not been successful to date^7,8^. Antitumor function of CAR T cells in solid tumors relies on efficient trafficking within the tumor microenvironment and sustained cytotoxic activity^9-12^. Since these processes involve dynamic cell-cell interactions, a comprehensive profiling of tumor-CAR T cell crosstalk and antigen-specific cytotoxicity are crucial to understanding mechanisms that enhance CAR T cell therapeutic efficacy.

Preclinical testing of CAR T cell cytotoxicity *in vivo* has provided valuable insights into identifying potent CAR T cell therapies^13,14^, however these models are costly and pose challenges for investigating dynamic cell-cell interactions. While intravital imaging allows for monitoring immune cell trafficking and cytotoxicity, its duration is limited to a few hours and control of the tumor microenvironment is challenging^15,16^. Several *in vitro* studies have employed cocultures of CAR T cells with tumor cells to measure CAR T cell function by colorimetric cytotoxicity assays^17-21^. While these assays offered a rapid and straightforward evaluation of CAR T cell function, they typically provide a single time-point measurement. Assays using micropatterning of tumor cell islands allowed for spatial segregation of each cell type and are compatible with time-lapse imaging^22^. However, as the tumor cells were plated on a two-dimensional surface, it is challenging to recreate the three-dimensional architecture of the solid tumor microenvironment.

Spheroids serve as more physiologically relevant 3D *in vitro* models, allowing assessment of drug sensitivity^23-26^ and tumor-immune cell signaling^27-29^. To model CAR T cell-cancer spheroid interactions, previous studies have used ultra-low adhesion well plates^17,30^ or hanging drop assays^31-33^. These methods are simple, requiring no special equipment, but they are labor-intensive and pose challenges in obtaining high-resolution images for the dynamic profiling of cell-cell interactions^34,35^. Microfabrication technologies present a promising approach to control and monitor T cell interactions with cancer spheroids. For example, microfluidic devices seeded with T cell receptor-engineered T cells and cancer spheroids revealed differences in T cell migratory behavior and cytotoxic efficiency between 2D and 3D *in vitro* environments^36^. Another microfluidic study demonstrated how enhanced vascularization of cancer spheroids is critical for CAR T cell delivery and cytotoxic function^37^. Furthermore, a microfluidic platform has previously shown the generation of cancer spheroids with co-encapsulation of stromal cells to test combination immunotherapies with CAR T cells^38^. Despite these advances, the spatiotemporal patterns of CAR T cell infiltration and cytotoxicity following sensing of a target antigen, inside cancer spheroids remain poorly understood.

In this study, we developed a microwell-based assay to monitor the dynamic processes of CAR T cell infiltration and cytotoxicity in 3D cancer spheroids. To evaluate the antigen-specific antitumor activity of CAR T cells, we utilized a switchable adaptor CAR system, SNAP-CAR, which mediates tumor antigen recognition through covalent attachment of a co-administered antibody adaptor bearing a benzylguanine motif^39^. This adaptor system allows for universal targeting of antigens and functional profiling within the same batch of engineered CAR T cells^40-50^. Specifically, we assessed CAR T cell cytotoxicity against HER2+ breast cancer spheroids in the presence or absence of an adaptor antibody that recognizes the HER2 antigen, Herceptin^51^. Through the fabrication of thin polydimethylsiloxane-based microwells using spin coating, we were able to perform high-resolution, live confocal imaging. Our microwell array facilitated monitoring of CAR T cell clustering and cytotoxicity in individual spheroids over time and led to the identification of the initial CAR T cell to spheroid area ratio as a predictor of cytotoxicity efficiency. Additionally, spatiotemporal analysis of CAR T cell functions, demonstrated distinct patterns of CAR T cell-mediated killing in the spheroid core compared to the periphery. These results demonstrate the utility of the microwell array platform for elucidating dynamic interactions of CAR T cells with 3D cancer spheroids following antigen sensing that promotes CAR T cell clustering and elimination of HER2+ breast cancer cells. Finally, our studies on CAR T cell function within the spheroid core have important implications for the design of cell-based immunotherapies to promote immune cell trafficking in solid tumors.

## Results

### Formation of 3D spheroids using HER2+ breast cancer cells in microwell arrays

We developed an array of microwells with controllable size to form 3D cancer spheroids under conditions that allowed high-resolution imaging. Microwells were fabricated using polydimethylsiloxane (PDMS) patterned onto a SU-8 silicon wafer using spin-coating to create devices with ∼270 µm thickness that are compatible with high numerical aperture objectives **(Fig.1a-c)**. The microwells were bonded onto a 24-well glass-bottom plate to enable stable long-term live cell microscopy and minimize evaporation of cell culture medium. Coating of microwell surfaces with a pluronic solution prevented cell adhesion and promoted cell-cell clustering under suspension culture. We found that HER2+ breast cancer cell lines BT474 and EFM192 could efficiently form spheroids in our microwell array **(Fig. 1c & Fig. S1)**. As expected, spheroid diameter increased for microwells with larger side lengths **(Fig.1d,e & Fig. S1a,b)**. We found that microwells with a side length of 50 µm included a spheroid that occupied the whole microwell area **(Fig.1f & Fig. S1c)**. Similarly, the majority of the microwell area was occupied with the spheroid for a side length of 100 μm. Finally, for microwells with a side length of 200μm, there was greater than 50-75% free surface area on the microwell, where another cell type could be introduced to monitor dynamic cell-cell interactions **(Fig.1f & Fig. S1c)**.

**Figure 1.**
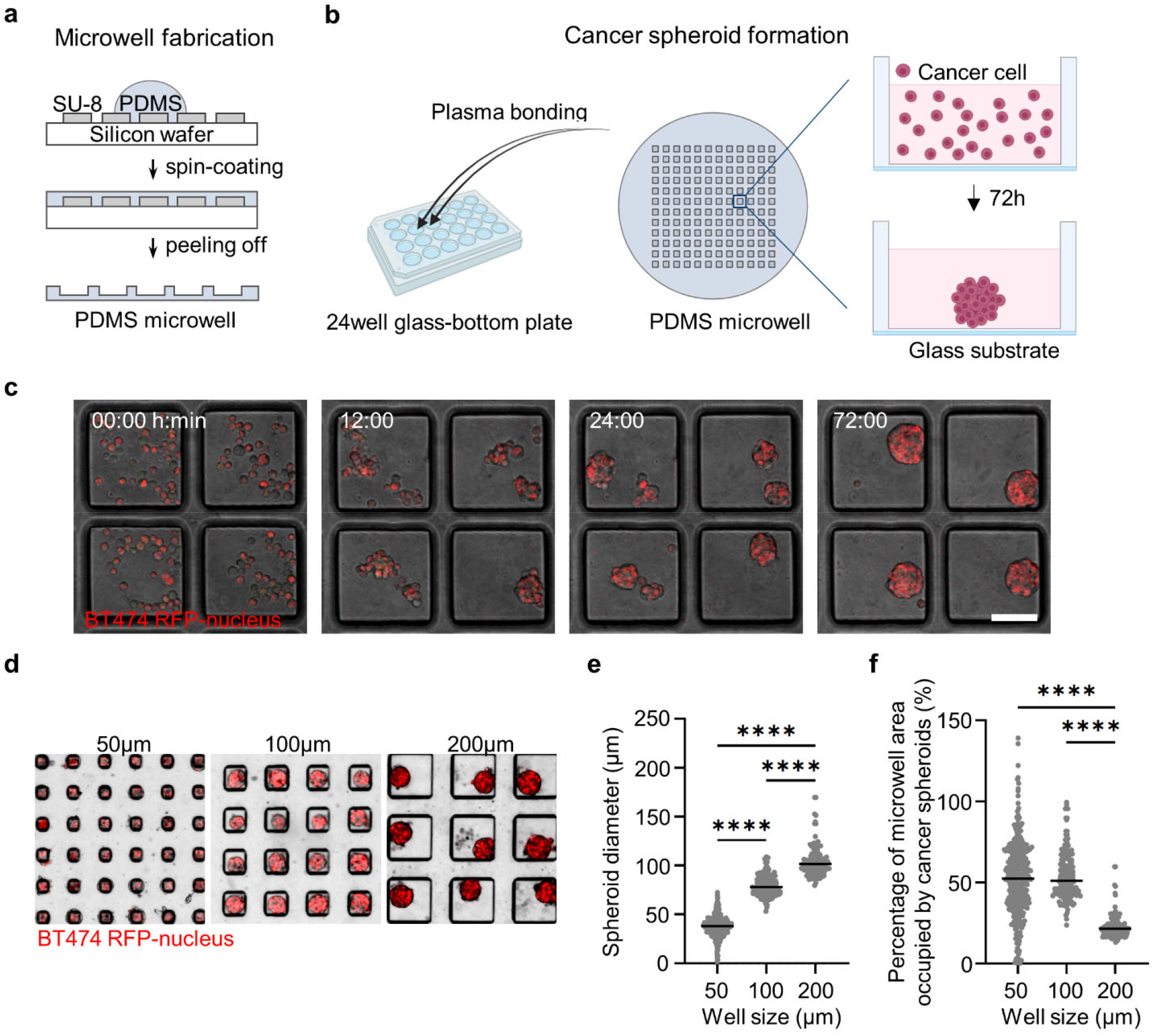
Microwell array platform to form and immobilize HER2+ breast cancer spheroids. (a) Schematics of PDMS microwell fabrication on SU-8 silicon wafer. (b) Bonding of PDMS microwell array on glass-bottom platform and cell seeding to form cancer spheroids. (c) Time-lapse images showing the spheroid formation of BT474-H2BRFP cells in microwells over 72 hrs. Scale bars, 100 µm. (d) Spheroid formation of BT474 cells in microwells with varying side length (50, 100, 200 µm). (e) Quantification of the diameter of BT474 spheroids formed in 50, 100, 200 µm microwells (N=392, 174, 119 respectively). (f) Quantification of the percentage of microwell area occupied by BT474 spheroids in microwells with different side lengths (50, 100, 200 µm). ^****^p<0.0001 (one-way ANOVA analysis).

### CAR T cell sensing of HER2 on cancer cells stimulates cytotoxicity and promotes CAR T cell infiltration within 3D cancer spheroids

To study antigen-specific, CAR T cell function with HER2+ breast cancer cells, we seeded CAR T cells in the microwells (200 μm x 200 μm) with cancer spheroids that were previously formed **(Fig. 2a)**. We utilized the SNAP-CAR “universal” adaptor CAR system, for which the SNAPtag protein takes the place of the antigen binding region, and instead of directly binding to an antigen on a cancer cell, covalently attaches to a benzylguanine (BG)-conjugated antibody^39^. Specifically, we used the HER2-specific antibody adaptor, BG-Herceptin^51^. This adaptor antibody binds to HER2+ cancer cells via the variable region of the antibody, while the BG motif reacts with the SNAPtag fusing it to the CAR T cells (**Fig. 2b**). To evaluate the performance of the adaptor-directed CAR T cells, we analyzed the overlap of a dead cell marker (cyan, **Fig. 2c**) with a cancer cell specific-marker (red, **Fig 2c & Fig S2**). We found higher cancer cell death for both BT474 spheroids and EFM192 spheroids in the presence of the BG-Herceptin adaptor compared to control cocultures with no adaptor. Specifically, 23% of cancer spheroid area colocalized with dead cell staining for BT474 spheroids treated with BG-Herceptin compared to 7% in the untreated control (p<0.0001) and a similar response was observed for EFM192 spheroids with 26% and 14% (p<0.0001), respectively (**Fig. 2d & Fig. S3**).

**Figure 2.**
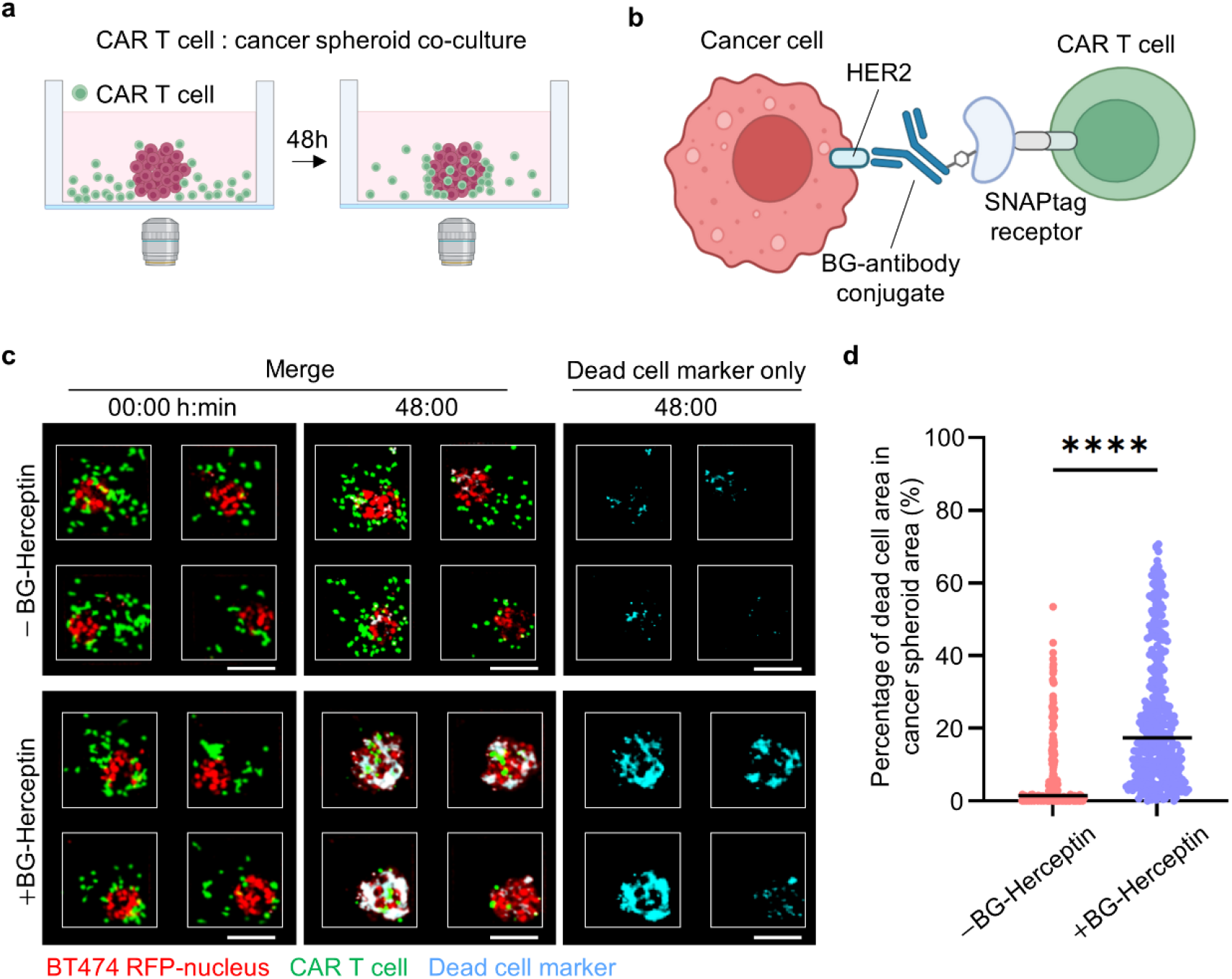
CAR T cell-mediated cytotoxicity in HER2+ breast cancer spheroids. (a) Setup of CAR T cell and cancer spheroid coculture. (b) Schematics of SNAP-CAR T cells interacting with BG-conjugated antibody adaptor to recognize HER2 antigen on cancer cells. (c) Images showing CAR T cell interactions with BT474 cancer spheroid s ± BG-Herceptin adaptor at 0 h and 48hrs. Red: BT474-H2BRFP, Green: CMFDA dye-stained CAR T cell, Cyan: Sytox deep red cell death dye. Scale bars, 100 µm. (d) Quantification of cytotoxicity levels, calculated as the overlap area between dead cell markers (Cyan) and spheroid (RFP) divided by the total spheroid area in BT474 cancer spheroids (N=192 microwells for –BG-Herceptin and N=377 microwells for +BG-Herceptin from 3 independent biological replicates) at 48 h. ^****^p<0.0001 (Unpaired t-test analysis).

Next, we used live cell confocal imaging to monitor the dynamic interactions of CAR T cells with BT474 cancer spheroids under both baseline (-BG-Herceptin) and antigen-sensing (+ BG-Herceptin) conditions (**Fig. 3a,b**). We found that following HER2-sensing the fraction of CAR T cells that interacted with cancer spheroids increased over time, with ∼73% of total CAR T cells interacting with cancer spheroids after 12 hrs (**Fig. 3c**). Under baseline conditions in the absence of BG-Herceptin, only ∼37% of total CAR T cells interacted with cancer spheroids after 12 hrs (**Fig. 3c**). Time-lapse imaging at a focal plane that captured the spheroid core, showed progressive infiltration of CAR T cells when treated with BG-Herceptin, while CAR T cells remained at the periphery of spheroids in the absence of BG-Herceptin treatment (**Fig. 3a, b**). To characterize the spatial patterns of CAR T cell trafficking within breast cancer spheroids, we divided each spheroid into three zones as a function of radius: (a) the spheroid periphery (outer zone), (b) the intermediate zone and (c) the spheroid core (inner zone) (**Fig. 3d**). We found that under baseline conditions (-BG-Herceptin), most of the CAR T cells (∼90%) remained at the spheroid periphery and there was no significant change of the CAR T cell population within the intermediate zone or spheroid core over time (**Fig. 3e**). However, following treatment with BG-Herceptin, the distribution of CAR T cells within the spheroid changed over time (**Fig. 3f**). Notably, 24 hrs following antigen sensing, a similar population of CAR T cells were present in the periphery (44%) and intermediate (39%) zones, with the lowest population being in the spheroid core (17%) (**Fig. 3f**).

**Figure 3.**
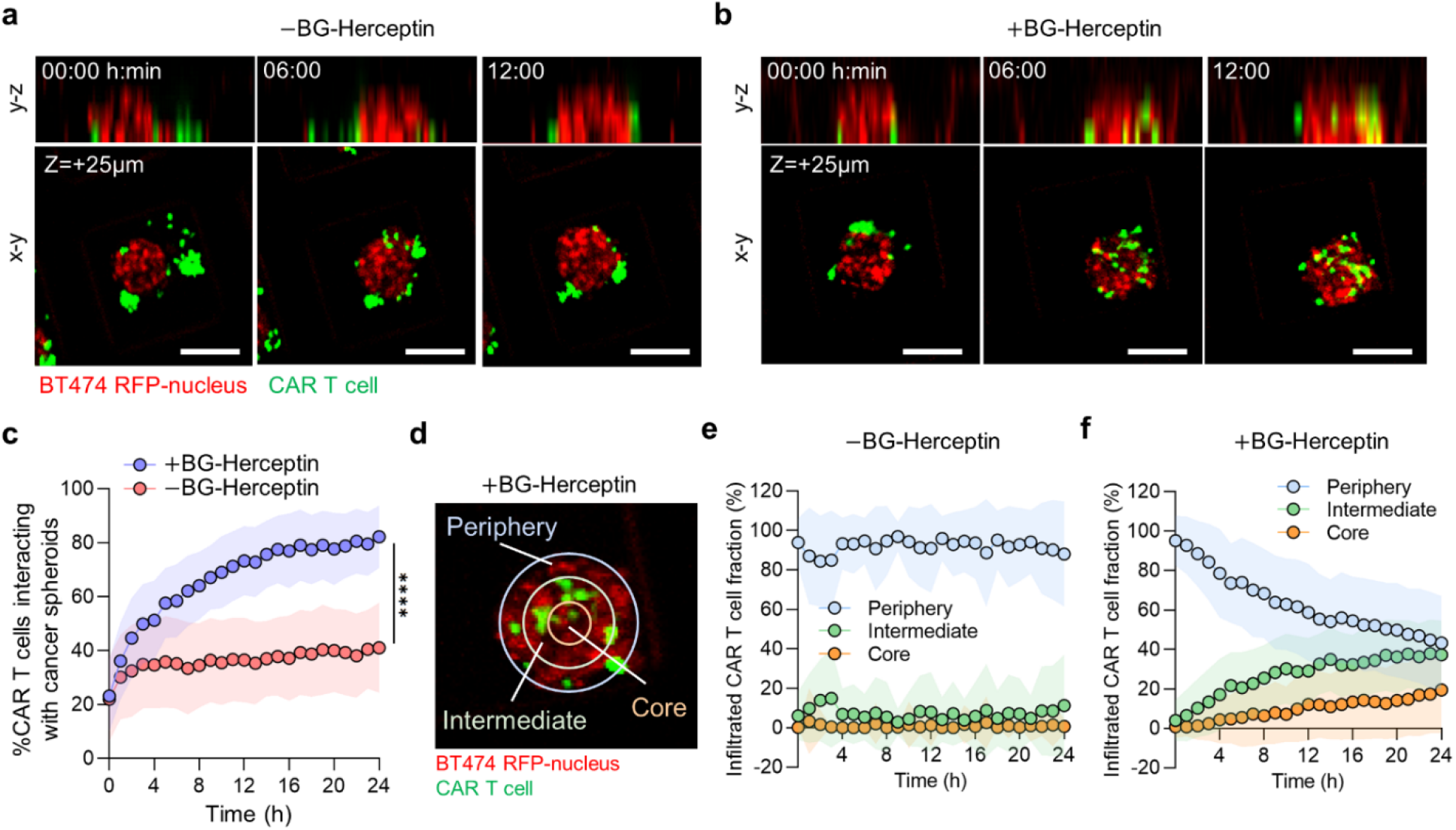
Sensing of HER2 on breast cancer cells by CAR T cells promotes CAR T cell infiltration within cancer spheroids. (a,b) Time-lapse confocal fluorescence imaging of y-z (top view) and x-y (side view) planes of CAR T cell-BT474 spheroid cocultures (a) in the absence and (b) presence of BG-Herceptin (HER2 sensing). Red: BT474 H2B-RFP, green: CMFDA dye-stained CAR T cells. x-y plane images are captured at z=+25 µm from microwell bottom surface. Scale bars, 100 µm. (c) Sensing of HER2 increases the percentage of CAR T cells interacting with cancer spheroids. The % interaction CAR T cells was calculated as the GFP area (CAR T cells) overlapping with RFP area (cancer cells) normalized by total GFP area within a microwell. Plots represent mean ± SD (N=145 microwells (+BG-Herceptin) and N=91 microwells (–BG-Herceptin). ^****^p<0.0001 (Unpaired t-test analysis). (d) Spatial analysis of CAR T cell infiltration in cancer spheroids divided in three zones as a function of radial position. To define the periphery and intermediate zones we set the difference between the inner and outer radii at 20μm for each zone, while the core occupied the remaining area of each spheroid. (e,f) Spatial distribution of infiltrated CAR T cells over time (e) in the absence of BG-Herceptin and (f) in presence of BG-Herceptin. Plots represent mean ± SD (N=145 microwells (+BG-Herceptin) and N=91 microwells (–BG-Herceptin). Results are representative of three biological replicates.

Given previous reports on CAR T cell clustering in two dimensional assays^22^, we assessed the clustering patterns of CAR T cells that had infiltrated within the spheroids and the impact of antigen-sensing. Infiltrated CAR T cells within cancer spheroids were classified as single cells (blue in **Fig. 4a**) or clusters (white in **Fig. 4a**). We found that following 24 hrs of HER2-sensing (+BG-Herceptin), 13% of infiltrated CAR T cells formed clusters, while only 0.5% of CAR T cells formed clusters under baseline conditions (-BG-Herceptin) (**Fig. 4b**). Analysis of CAR T cell cluster area within cancer spheroids over time showed a rapid increase following treatment with BG-Herceptin (**Fig. 4c**). Taken together, these results show that sensing of the HER2 antigen promotes CAR T cell-mediated killing of breast cancer cells and stimulates CAR T cell infiltration and cluster formation within cancer spheroids.

**Figure 4.**
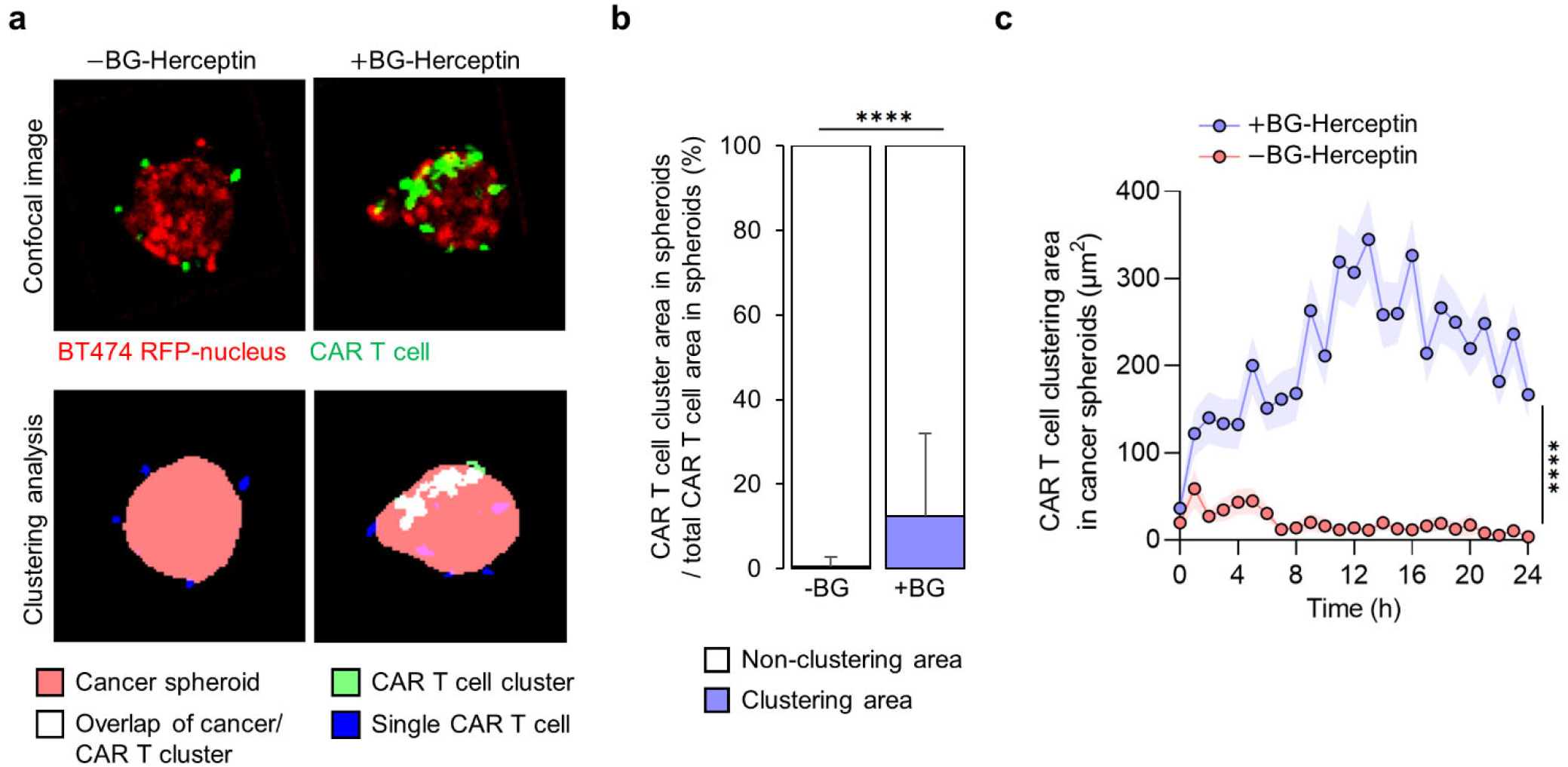
CAR T cell clustering within breast cancer spheroids depends on HER2 sensing. (a) Image analysis of CAR T cell clustering in cancer spheroids at z=+25 µm focal plane in the absence and presence of BG-Herceptin (HER2 sensing). Images in top row are fluorescence images and images in the bottom row are segmentations based on cancer spheroid area (red), single CAR T cell area (blue), CAR T cell cluster area (green), and CAR T cell cluster area overlapped with cancer spheroid area (white). (b) Percentage of CAR T cell clustering area over total CAR T cell area in cancer spheroids at 24 h ± BG-Herceptin. Mean ± SD. N=145 microwells (+BG-Herceptin) and N=91 microwells (–BG-Herceptin). (c) The size of CAR T cell cluster area within cancer spheroids increases over time only when HER2 is sensed. Mean ± sem (N=145 microwells (+BG-Herceptin) and N=91 microwells (– BG-Herceptin). Results are representative of three biological replicates. ^****^p<0.0001 (Unpaired t-test analysis).

### Dynamic profiling at the individual spheroid level uncovers predictors of CAR T cell-mediated cytotoxicity

Next, we characterized the cytotoxic activity kinetics of CAR T cells that infiltrated within cancer spheroids. Consistent with our end-point results, we found that BT474 cancer spheroids treated with the BG-Herceptin adaptor exhibited higher levels of cancer cell death for all timepoints compared to the no adaptor condition, with killing initiating 10 hrs following HER2 sensing (**Fig. 5a,b**). Specifically, under treatment with BG-Herceptin CAR T cells showed an increased rate of antitumor cytotoxicity over time (0.59% of tumor cell area killed per hr between 10-48hrs), compared to the baseline cytotoxicity in the absence of antigen sensing (0.28% of tumor cell area killed per hr between 10-48hrs. p<0.0001) (**Fig. 5b & Fig. S4**). Furthermore, this antigen-specific CAR T cell-mediated death was significantly higher than the cancer spheroid only control (6.2 fold, p<0.0001) (**Fig. 5b**).

**Figure 5.**
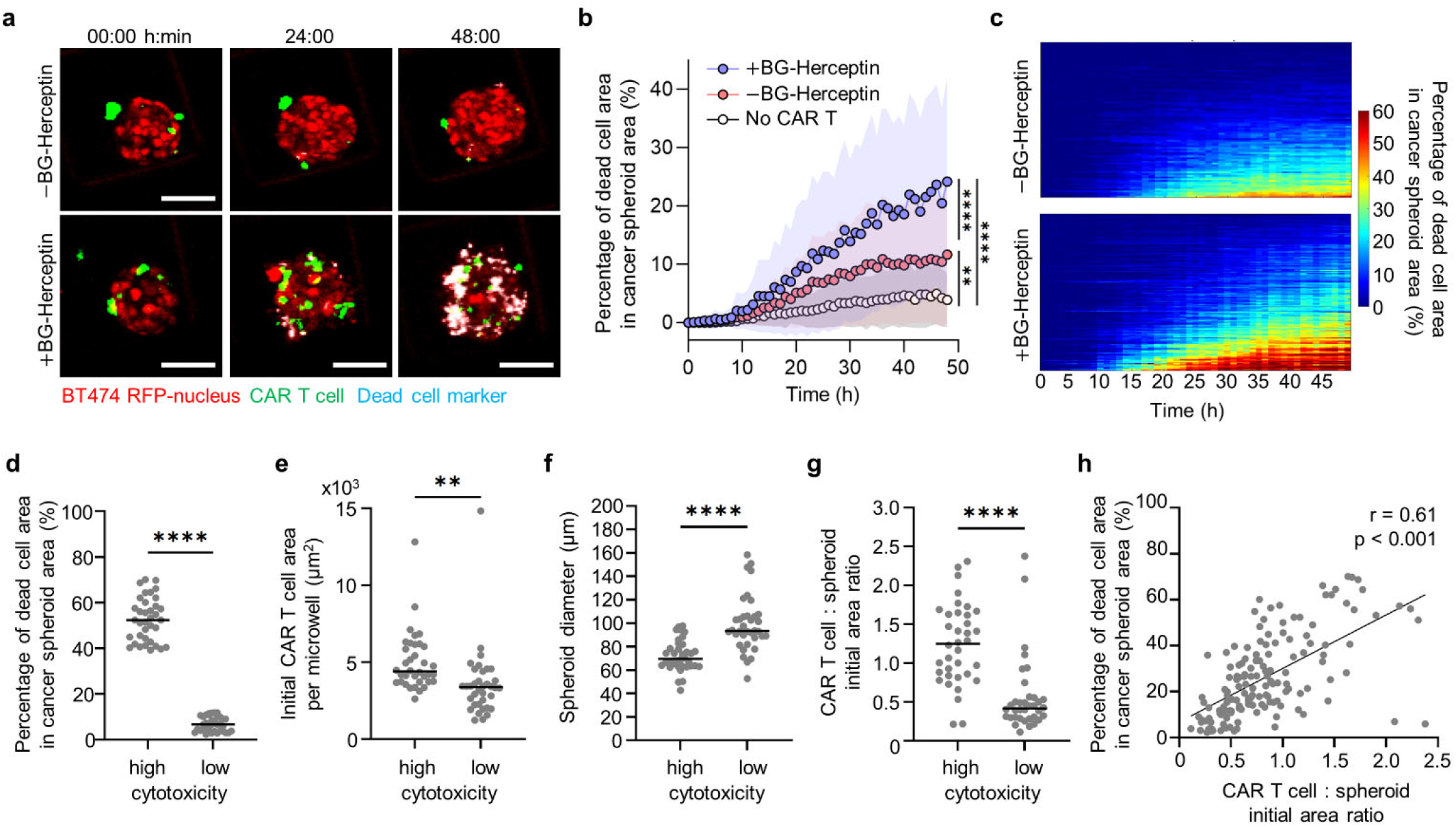
Dynamic profiling of CAR T cell cytotoxic killing reveals initial CAR T : spheroid ratio as a predictor of antitumor cytotoxicity. (a) Time-lapse confocal fluorescence images of CAR T cell - BT474 spheroid coculture at z=+25 µm in the absence and presence of BG-Herceptin. Red: BT474-H2BRFP, Green: CMFDA dye-stained CAR T cells, Cyan: dead cell marker. Scale bars, 100 µm. (b) Quantification of cytotoxicity levels in BT474 cancer spheroids over time ± BG-Herceptin. Mean ± SD (-BG: N=91; +BG: N=145; no CAR T: N=9 wells). (c) Heatmaps showing the temporal evolution of cytotoxicity for individual spheroids ± BG-Herceptin (Data from 91 microwells (-BG) and 145 microwells (+BG)). Each row in x-axis represents time and columns in y-axis represent individual microwells. (d-g) Each dot is a spheroid and two groups are shown based on high (top quartile) and low (bottom quartile) cytotoxicity levels from +BG-Herceptin condition in panel c. Characterization of (d) cytotoxicity, (e) initial CAR T cell area, (f) initial spheroid diameter, and (g) initial CAR T cell : spheroid area ratio. ^**^p<0.01, ^****^p<0.0001 (Unpaired t-tests). (h) Correlation between cytotoxicity and initial CAR T cell : spheroid area ratio in microwells treated with BG-Herceptin (N=145, Pearson correlation coefficient r = 0.61, p<0.001).

We also evaluated CAR T cell cytotoxic killing at the individual spheroid level. There was significant heterogeneity in the cytotoxic efficacy with a range from 2 to 70% of spheroid area that was positive for dead cell marker at 48hrs (**Fig. 5c**). Furthermore, the temporal evolution patterns across spheroids were heterogeneous (**Fig. 5c**). To identify factors that drive these heterogeneous cytotoxic outcomes, we first compared the microwells with the highest (top quartile with 53±7% of spheroid area overlapping with dead cell marker) and lowest cytotoxicity (bottom quartile with 10±3% of spheroid area overlapping with dead cell markers **Fig. 5d**). We found that the top quartile of microwells with high cytotoxicity exhibited a higher number of initial CAR T cells compared to the bottom quartile (**Fig. 5e**). Furthermore, the initial spheroid diameter was significantly larger in the bottom quartile microwells, indicating that cancer cells in larger spheroids are less effectively killed (**Fig. 5f**). To account for both parameters, we calculated the initial ratio between CAR T cell and spheroid area which also effectively discriminated between the top quartile of microwells with high cytotoxicity compared to the bottom quartile of microwells with low cytotoxicity (CAR T cell : spheroid area ratio was 1.16±0.55 for top vs 0.48±0.38 for bottom quartiles, p<0.0001, **Fig. 5g)**. This relationship was also supported by evaluating the initial CAR T cell : spheroid area ratio and cytotoxicity outcomes across all spheroids (Pearson correlation coefficient r = 0.61, p<0.001) (**Fig. 5h**). We also found a positive correlation between a high initial CAR T cell : spheroid area ratio and high cytotoxicity in the EFM192 breast cancer spheroids (**Fig. S5**).

### Spatiotemporal analysis of CAR T cell cytotoxicity in the spheroid periphery and core

We next investigated the spatial patterns of cytotoxic CAR T cell activity within individual spheroids, while increasing the CAR T cell seeding density in the microwell (from 0.2E6 cells/ml to 0.8E6 cells/ml, **Fig 6a, Fig S6a, b**). In microwells seeded with a higher number of CAR T cells, we found a higher number of CAR T cells within the spheroid periphery for both treatment with BG-Herceptin and the baseline conditions (**Fig 6b**). However, CAR T cell recruitment within the spheroid core increased only when BG-Herceptin was added to facilitate sensing of the HER2 antigen (**Fig 6c**). Analysis of CAR T cell-mediated cytotoxicity in the spheroid core compared to the periphery revealed that in the spheroid periphery cancer killing was observed at 10hr, while killing in the spheroid core was delayed to 20hr (**Fig 6d, e**). At the higher CAR T cell seeding densities under BG-Herceptin treatment, the killing rates at the periphery (0.97%/hr between 10-48hrs) and core (0.94%/hr between 20-48hrs) were comparable (**Fig 6d, e**). However, at lower CAR T cell seeding densities, the killing rate at the periphery (0.58%/hr between 10-48hrs) was higher than the killing rate at the core (0.39%/hr between 20-48hrs). Furthermore, consistent with our analysis of cytotoxicity predictors (**Fig 5h**), we found that increasing the CAR T cell seeding density within the microwell resulted in a higher initial CAR T cell : spheroid area ratio (1.9±0.61) that was also associated with higher cytotoxicity (∼50%) **(Fig S6b)**. These results show that antigen-specific CAR T cell : tumor cell interactions promote CAR T cell infiltration within a 3D spheroid core and subsequent tumor cell killing.

**Figure 6.**
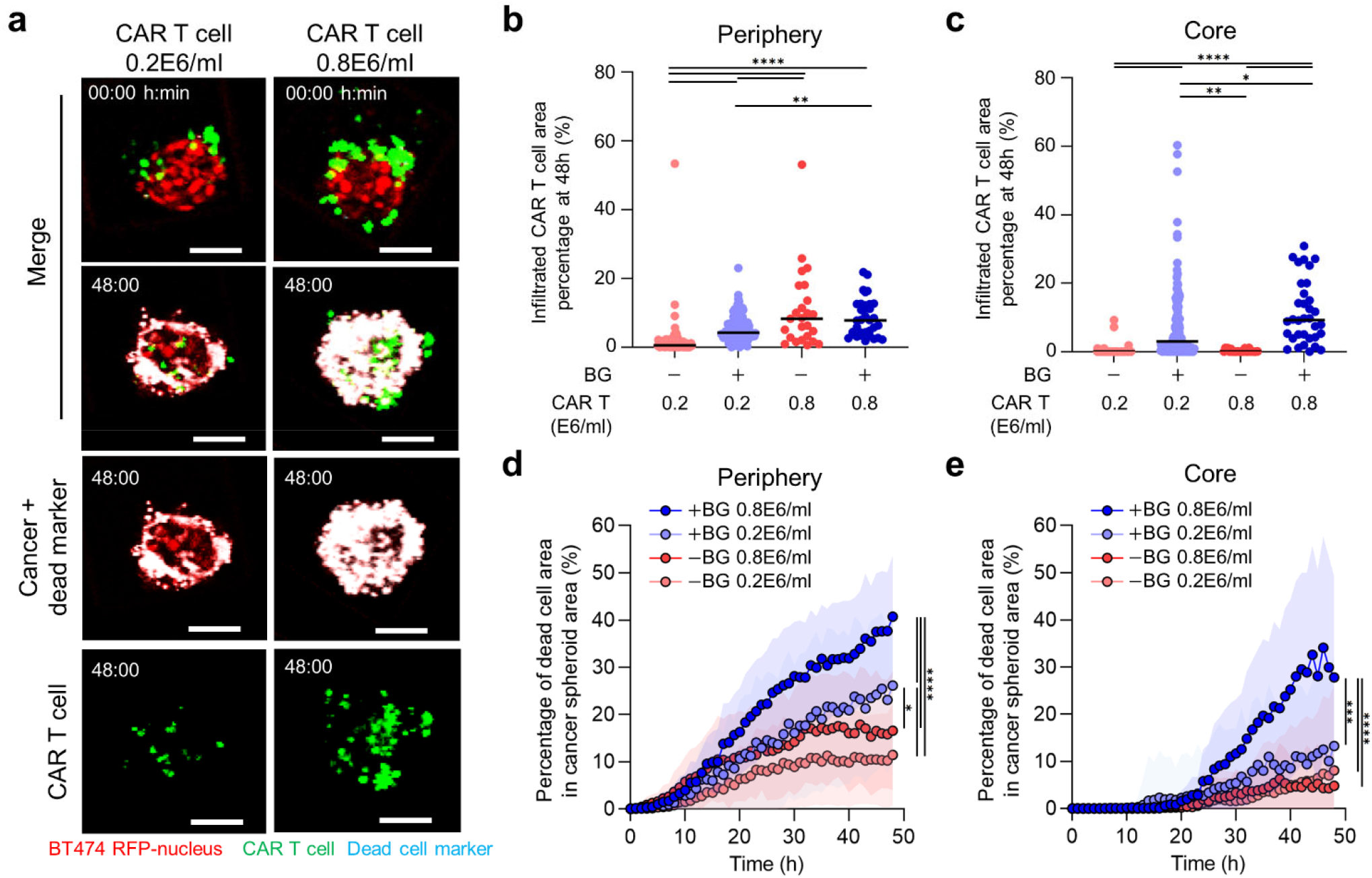
CAR T cell cytotoxicity within the spheroid core is enhanced by higher CAR T cell density when HER2 antigen is recognized. (a) Time-lapse confocal fluorescence imaging of CAR T cell-BT474 spheroid coculture in the presence of BG-Herceptin with different seeding densities (low: 0.2E6 cells/ml and high: 0.8E6 cells/ml) at 0 h and 48hrs. Red: BT474-H2BRFP, Green: CMFDA dye-stained CAR T cells, Cyan: dead cell marker. Scale bars, 100 µm. (b,c) Quantification of % infiltrated CAR T cells out of total CAR T cells in each spheroids at (b) the periphery and (c) in the core at 48hrs ± BG-Herceptin with different CAR T cell seeding densities. (d,e) Cytotoxic activity (% spheroid area overlapping with dead cell markers) in (d) spheroid periphery and (e) core ± BG-Herceptin. Plots represent mean ± SD (-BG low (N=91), +BG low (N=145), -BG high (N=25), +BG high (N=34). ^*^p<0.05, ^**^p<0.01, ^***^p<0.001 ^****^p<0.0001 (one-way ANOVA analysis).

## Discussion

The solid tumor microenvironment poses a significant barrier that CAR T cells have to overcome in order to detect and kill cancer cells. Experimental models that faithfully recapitulate solid tumor architecture are needed to understand CAR T cell trafficking and test their antitumor efficacy. Here, we developed a microwell-based platform to quantitatively assess CAR T cell infiltration and cytotoxic functions in 3D HER2+ breast cancer spheroids. By profiling individual cancer spheroids, we identified the initial CAR T cell : spheroid area ratio as a predictor of CAR T cell-mediated killing. Furthermore, using spatiotemporal analysis of CAR T cell trafficking, we showed that HER2 antigen-sensing promotes CAR T cell clustering and infiltration within the 3D spheroid core and further enhances cytotoxicity. These results demonstrate the capabilities of our microwell assay as a miniaturized technology to elucidate dynamic cell-cell interactions at play during CAR T cell anti-tumor functions.

Spheroids enable the study of physiologically relevant parameters, including spatial zonation and cell motility within a multicellular environment, which is not feasible in 2D models^52^. Studies using cancer spheroids have yielded insights into drug distribution as a function of distance from the spheroid core^34,53^ and heterotypic tumor-immune cell interactions^54,55^. Compared to traditional suspension culture methods (e.g. hanging drop and ultra-low adhesion surfaces), integrating spheroids within microfabricated platforms provides advantages with respect to controlling cell seeding and the extracellular environment^56^. For example, microfluidic devices with cancer spheroids embedded in a 3D matrix facilitated analysis of directional T cell trafficking and cytotoxic activity^36,37,57^. Our microwell platform offers several critical features for monitoring tumor-immune cell crosstalk, in real-time, within a 3D cancer spheroid. First, spheroids are spatially segregated to facilitate parallel comparisons between a large number of wells in a single experiment. Second, 3D spheroids are formed in a microwell with controlled size. Finally, by controlling the cell seeding sequence, we avoid transferring cancer spheroids between culture platforms prior to high-resolution imaging.

Cell migration in a complex 3D microenvironment^58,59^ is critical for engagement of cytotoxic T cells with target tumor cells^36,60^. We utilized universal adaptor CAR T cells to study antigen-specific effects on migration under control of an adaptor molecule. We found that CAR T cell infiltration and clustering within cancer spheroids was enhanced following sensing of HER2 on cancer cells. Our results are consistent with previous *in vitro* studies that reported clustering under 2D conditions when comparing mock-transduced CAR T cells with anti-BCMA (B-cell maturation antigen) targeted CAR T cells^22^. Another study that employed 3D macroscale cultures also demonstrated that engagement of cognate targets by cytotoxic T cells promotes swarming^57^. These cooperative interactions between T cells following antigen recognition were also shown using a microfluidic droplet assay with melanoma spheroids and ovalbumin-targeted murine OT-I cytotoxic T cells^57^. Furthermore, we found that CAR T cells localized within the spheroid core only in the presence of HER2 antibody adaptor and irrespective of the CAR T cell seeding density. These results provide further supportive evidence for the critical role of antigen-specific tumor-immune crosstalk that mediates CAR T cell trafficking in a 3D environment.

Sustained CAR T cell cytotoxic activity represents a major challenge in developing CAR T cell therapies against solid tumors^61,62^. By monitoring CAR T cell-mediated tumor cell killing at the individual spheroid level, we showed the suppressive effects of spheroid size and that the initial CAR T cell to spheroid area ratio can serve as a predictor of cytotoxicity. These results agree with a previous study that showed killing of B16 melanoma cells expressing SIINFEKL as a function of initial OT-I T cell concentration^63^. Another study employed intravital imaging and demonstrated that a high local density of OT-I cytotoxic T cells correlated with a lower number of virus-infected cells *in vivo*^64^. It is also important to monitor cytotoxic T cell function in spatial zones that have been associated with poor T cell infiltration, such as the intratumor core^65,66^. Using spatiotemporal analysis of killing dynamics in our microwell platform, we showed that HER2+ cancer cells within the spheroid core could be eliminated by CAR T cells in an antigen specific manner. Our results are consistent with a previous study that formed HER2 therapy-resistant JIMT1 spheroids using ultra-low adhesion 96-well plates and evaluated cytotoxicity using anti-HER2 CAR T cells^31^.

In summary, we present a microwell platform to quantitatively study CAR T cell infiltration and cytotoxicity in HER2 breast cancer spheroids. Our findings reveal distinct patterns of anti-tumor T cell functions within the spheroid periphery compared to the spheroid core that are dependent on engagement of the HER2 antigen on tumor cells. The universal CAR T cell system used in our study provides flexibility in directing CAR T cells against antigens that can be targeted using a benzylguanine-conjugated antibody. Finally, our multiwell plate-compatible assay can be integrated into combinatorial drug screening platforms and the miniaturized scale offers advantages for studying low-volume, patient-derived samples.

## Materials and methods

### PDMS microwell fabrication and assembly

SU-8 silicon wafers served as molds for PDMS microwell fabrication. PDMS was mixed in a 10:1 ratio of elastomer base to curing agent and placed in a vacuum desiccator for one hour to degas. Then, 2.5g of the PDMS mixture was poured on the center of the microwell mold. The mold was fixed on a spin-coater and rotated at 1000 rpm for 5 minutes. Next, the mold was incubated overnight in an 80°C oven. Subsequently, the thin PDMS microwell sheet was peeled off and plasma-bonded into each well of a 24-well glass-bottom plate. After incubating the plate in an 80°C oven for 10 minutes, the plate was treated with plasma once more, and 500 µl of 2% pluronic solution was added to each well. The plate was then centrifuged at 900 rpm for 5 minutes to remove any bubbles trapped in the microwells. After incubating the plate in the incubator for one hour, it was washed three times with PBS and once with cell culture medium before seeding the cells.

### 3D spheroid formation and CAR T cell transduction

The human breast cancer cell lines BT474 and EFM192 (expressing fluorescent nuclear marker H2B-RFP^67^) were cultured in RPMI supplemented with 10% Heat Inactivated Fetal Bovine Serum (HIFBS) and 1% penicillin/streptomycin. A cancer cell seeding solution at a density of 0.2E6 cells per ml was prepared and subsequently 1 ml of the cancer cell solution was added to each well of a 24-well glass-bottom plate. The plate was centrifuged at 900 rpm for 1 minute to promote cancer cell settling within the microwells. We allowed 72 hrs for spheroids to form in a 5% CO2 and 37C incubator.

CAR T cells were generated by following a protocol described in our previous studies^39^. In brief, CD3+ T cells isolated with Pan T cell isolation kit (Miltenyi Biotec), were stimulated with TransAct Human T cell activation reagent (Miltenyi Biotec), 100 U/ml human IL-2 IS (Miltenyi Biotec) and 1 ng/ml IL-15 (Miltenyi Biotec) for 48 hrs. For lentiviral transduction, the viral vector-containing medium was spun down in retronectin coated plate at 2000 x g for 2 hrs at 32 °C, and after removing 2 ml of supernatant, 1 × 10^6^ of T cells were added in 4ml of media and spun down for an additional 10 mins 1000 x g at 32 °C. Cells were then expanded every 2-3 days, and fresh IL-2 and IL-15 were added in each expansion time. Transduction efficiency was tested at day 8 post-transduction (**Fig S7**).

### Production of BG-Herceptin

The anti-HER2 antibody trastuzumab (Herceptin, Genentech) was conjugated to BG following the protocol described in our previous studies^39^. In brief, Herceptin was buffer exchanged in PBS by using 7 K MWCO Zeba Spin Desalting Columns (ThermoFisher Scientific). Then it was incubated with 20ME (Molar equivalent) of BG-GLA-NHS (NEB) for 30 minutes at room temperature, subsequently buffer exchanged by using 7 K MWCO Zeba Spin Desalting Columns and the concentration was measured by Nanodrop One (Thermofisher Scientific).

### CAR T cell - spheroid interaction assays

Before seeding CAR T cells, we carefully aspirated the media from the microwells containing cancer spheroids and added 500 µl of RPMI media supplemented with a 1:5000 dilution of SYTOX™ Deep Red Nucleic Acid Stain dye (#S11381, Invitrogen) for tracking dead cells. To quantify CAR T cell numbers, we performed staining with 10 ng/ml CellTracker Green CMFDA Dye (#C7025, Fisher) for 25 minutes at 37°C and washed once with RPMI. Subsequently, we added 0.2E6 CAR T cells in RPMI supplemented with BG-Herceptin to each microwell at a final concentration of 1 μg/ml for +BG-Herceptin samples. For the higher initial CAR T cell seeding density, we added a four times higher concentration at 0.8E6 CAR T cells to the microwells. The cancer spheroid-only condition (no CAR T cells added) also included a concentration of 1 μg/ml of BG-Herceptin.

### Time-lapse confocal microscopy

For time-lapse imaging, the microwell plate was mounted on a Zeiss LSM700 confocal microscope housed within the Tokai-Heat incubation system. Live images were captured as a z-stack (3 stacks with a 25 µm interval starting from the bottom of the microwell surface) every hour for 48 hours using a 10X 0.45NA objective lens.

### Image analysis

MATLAB was utilized for all image analysis. To allow for whole cancer cell spheroid segmentation, we applied a 20 pixel median filter on the RFP (cancer cells) images to effectively group the nuclei of cancer cells into a single object prior to intensity thresholding. GFP (CAR T cells) and Cy5 (dead cell marker) images were also segmented through intensity thresholding. The percentage of CAR T cells interacting with cancer spheroids was determined by calculating the overlap area between the cancer spheroid area (RFP) and the CAR T cell area (GFP), divided by the total CAR T cell area (GFP) per microwell. The total CAR T cell area was computed by summing the GFP area within the microwell over the entire z-stacks. The cytotoxic metric in each cancer spheroid was determined by calculating the overlap area between the cancer spheroid area and dead cell area, divided by the total cancer spheroid area. To characterize CAR T cell clustering areas, we first identified GFP objects with an area above 50 pixels (312.5 µm^2^), comprising approximately ≥ 4 cells, as a CAR T cell cluster. The remaining GFP objects were classified as single CAR T cells. We also report the percentage of CAR T cells that were organized in a cluster by calculating the ratio of the overlap between the cancer spheroid area and the CAR T cluster area, divided by the overlap between the cancer spheroid area and total CAR T cell area.

### Statistical Analysis

For statistical analyses involving multiple groups, we applied one-way analysis of variance (ANOVA) and Tukey’s post hoc test for multiple comparisons. Two group comparisons were conducted using an unpaired t-test using Prism. The statistical significance is marked by asterisks in the figures, and we considered P-values below 0.05 as significant.

## Data availability

Data and computational code are available from the corresponding author upon request.

## Acknowledgements

This work was supported by the National Cancer Institute (R00CA222554 to I.K.Z), National Cancer Center Postdoctoral Fellowship (to Y.C), NIH grant R01 GM142007 (to J.L.), a Magee Women’s Research Institute Breast Cancer pilot grant, the UPMC Hillman Cancer Center and the Department of Bioengineering, Swanson School of Engineering at the University of Pittsburgh.

## Author contributions

Y.C, E.R, J.L and I.K.Z designed research; Y.C, M.L, T.B, R.L performed experiments; E.R and J.L generated SNAP-CAR T cells; Y.C, M.L, T.B analyzed data; Y.C, E.R, J.L and I.K.Z discussed data; Y.C, I.K.Z wrote the paper with input from all authors.

