## Supplementary Information for "CAR T cell infiltration and cytotoxic killing within the core of 3D breast cancer spheroids under control of antigen sensing in microwell arrays"

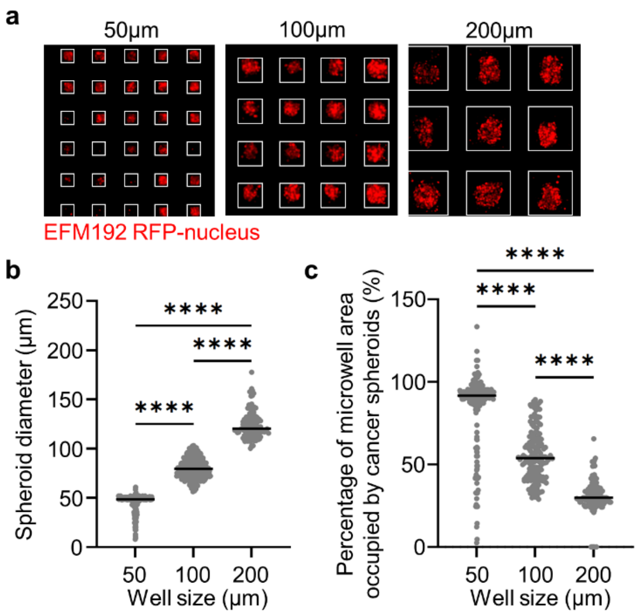


**Figure S1.** (a) Spheroid formation of EFM192 cells in microwells with varying side length (50, 100, 200 µm). (b) Quantification of EFM192 spheroid diameters in microwells with different size (N= 272, 437, 113, respectively). (c) Quantification of the percentage of microwell area occupied by EFM192 spheroids in each microwell design. ****p<0.0001.

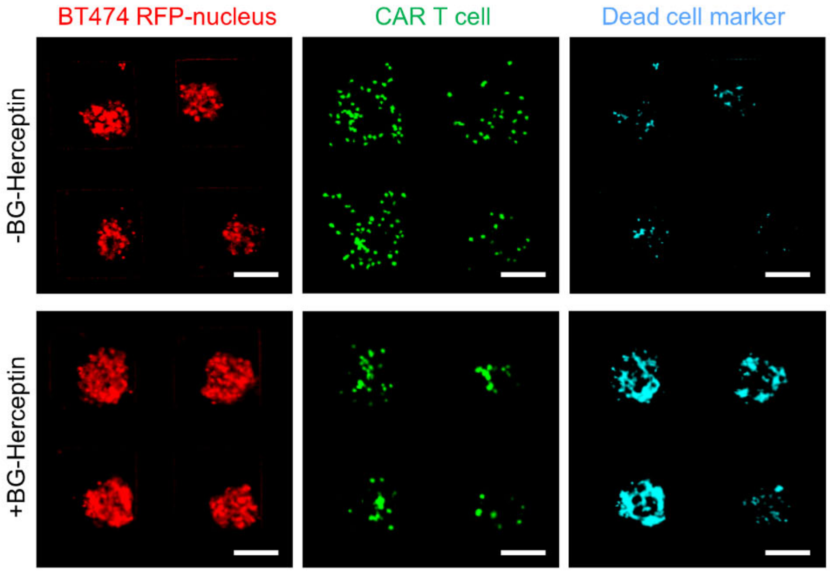


**Figure S2.** Separate fluorescent channel images of Fig 2C at 48 hrs of CAR T – BT474 spheroid co-cultures. Scale bars, 100 μm.

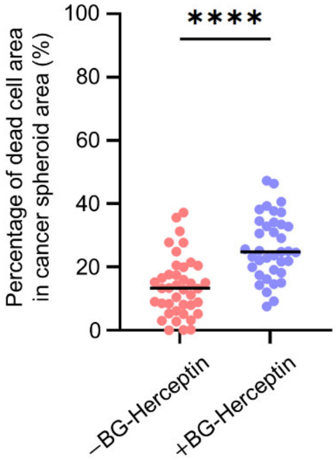


**Figure S3.** Quantification of cytotoxic levels, calculated at the ratio of dead cell area divided by total spheroid area for EFM192 cancer spheroids (N=40 microwells for –BG-Herceptin and N=37 microwells for +BG-Herceptin).

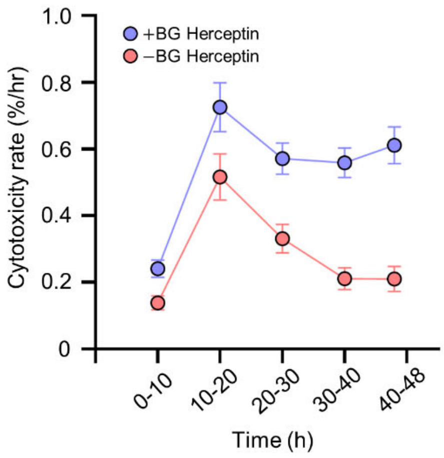


**Figure S4.** Cytotoxicity rate (calculated by diving the % cytotoxicity levels with each 10hr time-window) in the presence and absence of BG-Herceptin treatment in BT474 spheroids. Mean ± sem (N=91 (-BG), N=145 (+BG)).

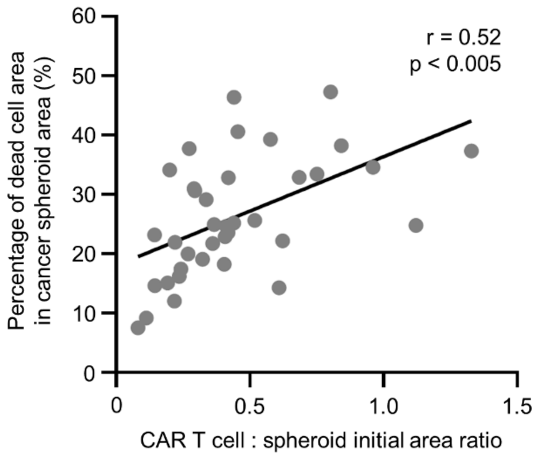


**Figure S5.** Correlation between cytotoxicity levels at 48 h and CAR T cell : EFM192 spheroid area ratio at t0 in microwells treated with BG-Herceptin (N=37, correlation coefficient $r = 0.52$, $p<0.005$).

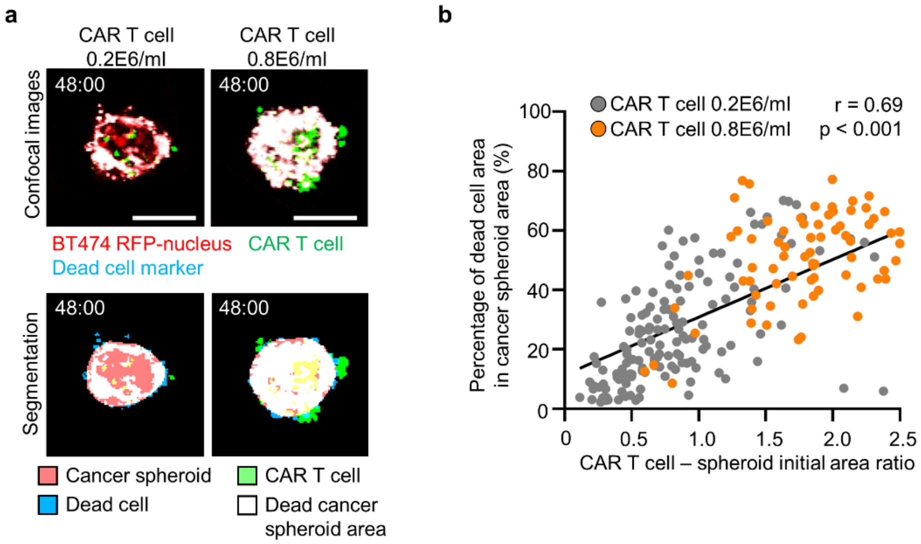


**Figure S6.** (a) Image analysis of CAR T cell infiltration and cytotoxicity in cancer spheroids at z=+25 μm focal plane with different CAR T cell seeding density (0.2E6 cell/ml and 0.8E6 cells/ml) at 0 h and 48 h. Images in a top row are confocal fluorescence images and images in bottom row are segmented images based on cancer spheroid area (red), CAR T cell area (green), dead cell area (cyan), and dead cell area overlapped with cancer spheroid area (white). (b) Correlation between cytotoxicity levels and CAR T cell : spheroid area ratio at t0 in microwells with different CAR T cell seeding density treated with BG-Herceptin in BT474 spheroids (correlation coefficient r = 0.69, p<0.001).

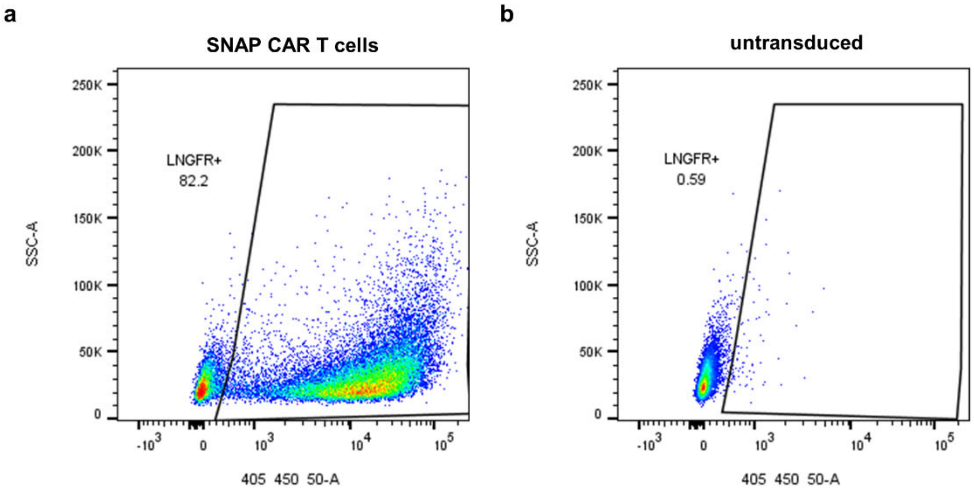


**Figure S7.** Flow cytometry analysis of the expression of SNAP-CAR on (a) transduced vs. (b) mock (untransduced) primary human T cells, assessed by recording LNGFR+ expression at day 8 post-transduction with the MSGV1-SNAP-41BBz-T2A-LNGFR retrovirus[1]. [antibody: anti-LNGFR BV421, clone C40-1457]"
